# Oncogenic Kras induces spatiotemporally specific tissue deformation through converting pulsatile into sustained ERK activation

**DOI:** 10.1101/2022.09.14.507992

**Authors:** Tianchi Xin, Sara Gallini, David Gonzalez, Lauren E. Gonzalez, Sergi Regot, Valentina Greco

## Abstract

Tissue regeneration and maintenance rely on coordinated stem cell behaviors. This orchestration can be impaired by oncogenic mutations leading to tissue architecture disruption and ultimately cancer formation. However, it is still largely unclear how oncogenes perturb stem cells’ functions to break tissue architecture. Here, we used intravital imaging and a novel signaling reporter to investigate the mechanisms by which oncogenic Kras mutation causes tissue disruption in the hair follicle. Through longitudinally tracking the same hair follicles in live mice, we found that KrasG12D, a mutation that can lead to squamous cell carcinoma, induces epithelial tissue deformation in a spatiotemporally specific manner. This tissue architecture abnormality is linked with a spatial dysregulation of stem cell proliferation as well as abnormal migration during hair follicle growth. By using a reporter mouse that allows us to capture real-time ERK signal dynamics at the single cell level, we discovered that KrasG12D, but not a closely related mutation HrasG12V, converts the pulsatile ERK signal fluctuation in the stem cells into sustained activation. Furthermore, by combining drug treatment with longitudinal imaging, we demonstrated that temporary inhibiting ERK signal reverts the KrasG12D-induced tissue deformation, suggesting that sustained ERK activation leads to tissue architecture disruption in Kras mutant hair follicles. Altogether, our work suggests that oncogenic mutations induce tissue abnormalities when spatiotemporally specific conditions are met, which allows mutant stem cells to disturb local cell coordination through altering dynamic signal communications.

## Introduction

Tissue function depends on the orchestration of cell organization and behaviors, which is mediated by dynamic cell-cell communication. The coordination of these events can be disrupted by oncogenic mutations, leading to various disorders such as cancer. However, the mechanisms by which oncogenic mutations initiate tissue disruption and cancer remain largely unknown due to the technical limitations of tracking the same cells as they transition from normal to oncogenic Furthermore, cancer formation is context dependent, as the cells with the same oncogenic mutations can lead to distinct consequences based on the specific microenvironments they reside in^1, 2^. Recent findings that phenotypically normal tissues can carry many oncogenic mutations^3^ further stress the importance of identifying the particular cancer-promoting contexts that allow tissue disruption to occur, which will ultimately allow us to understand the mechanisms of cancer initiation.

Skin epithelium is a powerful system for studying cancer biology because of its well-characterized cancer types and associated oncogenes, as well as the mouse models that recapitulate human cancers^4^. Cutaneous squamous cell carcinoma (cSCC) is one of the most frequent cancers originated from the skin epithelium. Mouse models with Ras oncogene expression in the skin epithelium have identified the population in the hair follicle that can initiate cSCC, as well as multiple types of tissue aberrancy before the typical cancer phenotypes appear^5–8^. However, the cellular and molecular alterations that induce those pre-cancer tissue abnormalities are still largely unclear, in part because the skin epithelium constantly regenerates and dynamically responds to oncogenes. For example, mutation-induced aberrancy in the skin epithelia can be corrected over time and the same oncogenic mutation can lead to distinct consequences in different epithelial compartments^9, 10^. These findings highlight the need to capture the spatiotemporal tissue dynamics along with the molecular and cellular changes that result from the acquisition of oncogenic mutations.

Among the skin epithelial tissues that are used to model cSCC, the hair follicle epithelium represents a powerful system for dissecting the context-dependent mechanisms of oncogenesis, as its cyclic nature creates diverse microenvironments for oncogenic mutant cells. First, as hair follicles go through different phases of the hair cycle, including rest (Telogen) and multiple stages of growth (Anagen) and regression (Catagen) phases, hair follicle epithelial cells receive drastically different signals from the mesenchyme and other niche components (Supplementary figure 1)^11^. Furthermore, during the stages of growth phase, hair follicle stem cells expand, reorganize and differentiate into multiple cell types to build a stereotypic architecture in which cells in different regions exhibit distinct cellular and molecular phenotypes^12–15^. For example, at the late growth stage, hair follicle epithelial cells form a structure of concentric layers, including the inner layers of multiple types of upward moving differentiating cells that are derived from the progenitors at the bottom of the hair follicle, as well as the outermost layer of expanded stem cells (Outer Root Sheath, or ORS cells) that proliferate and migrate to sustain the hair follicle growth and refill the progenitor pool. Notably, even within the same layer, cells behave differently based on their positions along the hair follicle. Particularly, the stem cells that undergo division are mostly restricted to the lower part of the late growth hair follicles^16^. Together, these characteristics make the hair follicle an excellent model system to interrogate how different niches can influence mutant cells’ behaviors through spatiotemporally specific cues and lead to particular consequences.

Here, by combining an inducible oncogenic Kras model with intravital imaging, we uncover new mechanisms of Ras mutation-induced tissue disruption. Longitudinal tracking of the same tissues in live mice showed that the KrasG12D mutation induces epithelial deformation in the hair follicle in a spatiotemporally specific manner. Through time lapse analyses and live extracellular signal-regulated kinase (ERK) reporter, we also find that the normal pulsatile ERK signals in the stem cells can be converted by KrasG12D into sustained activation, which is associated with abnormal cell division and migration. Furthermore, via partial inhibition of ERK activation, we demonstrate that sustained ERK signaling induced by KrasG12D is required for the formation and maintenance of the epithelial deformation. Together, our work suggests that spatiotemporally specific tissue disruption can be induced by oncogenic mutations through modulating signaling dynamics.

## Results

### KrasG12D induces tissue deformation in a spatiotemporally specific manner during hair follicle regeneration

Several months after being induced in hair follicle stem cells, oncogenic KrasG12D mutation can cause the formation of Papilloma as well as cSCC if combined with the loss of tumor suppressor genes^5–8^. Before tumor emergence, multiple types of architecture disruption were identified including follicular hyperplasia and cyst formation. However, it is still not clear when and how these tissue abnormalities arose with respect to their resident niches. Within the months prior to tumor formation, hair follicles could go through multiple cycles of regeneration, which dramatically changed the microenvironments experienced by the mutant cells.

In order to understand the effect of different niches on mutant cells’ ability to disrupt tissue architecture, we first used live imaging to longitudinally track hair follicles with KrasG12D mutant epithelial cells to identify when and where during the hair cycle mutant cells cause specific types of tissue disruption. A driver that can target most stem cells in the hair follicle (Lgr5-CreER) was used to induce KrasG12D expression (LoxP-STOP-LoxP-KrasG12D) in the mice that also express an epithelial nuclear marker (K14-H2BGFP), right before the first rest phase of the hair cycle (Figure 1a). The same skin of the live mice was then imaged every 3-4 days throughout the growth and regression phases (Figure 1b). By incorporating a Cre reporter (LoxP-STOP-LoxP-tdTomato), we estimated that the Kras mutant allele was induced in the vast majority of the hair follicle stem cells in the rest phase as well as their progeny during the following phases in the experimental mice (Figure 1c).

**Figure 1.**
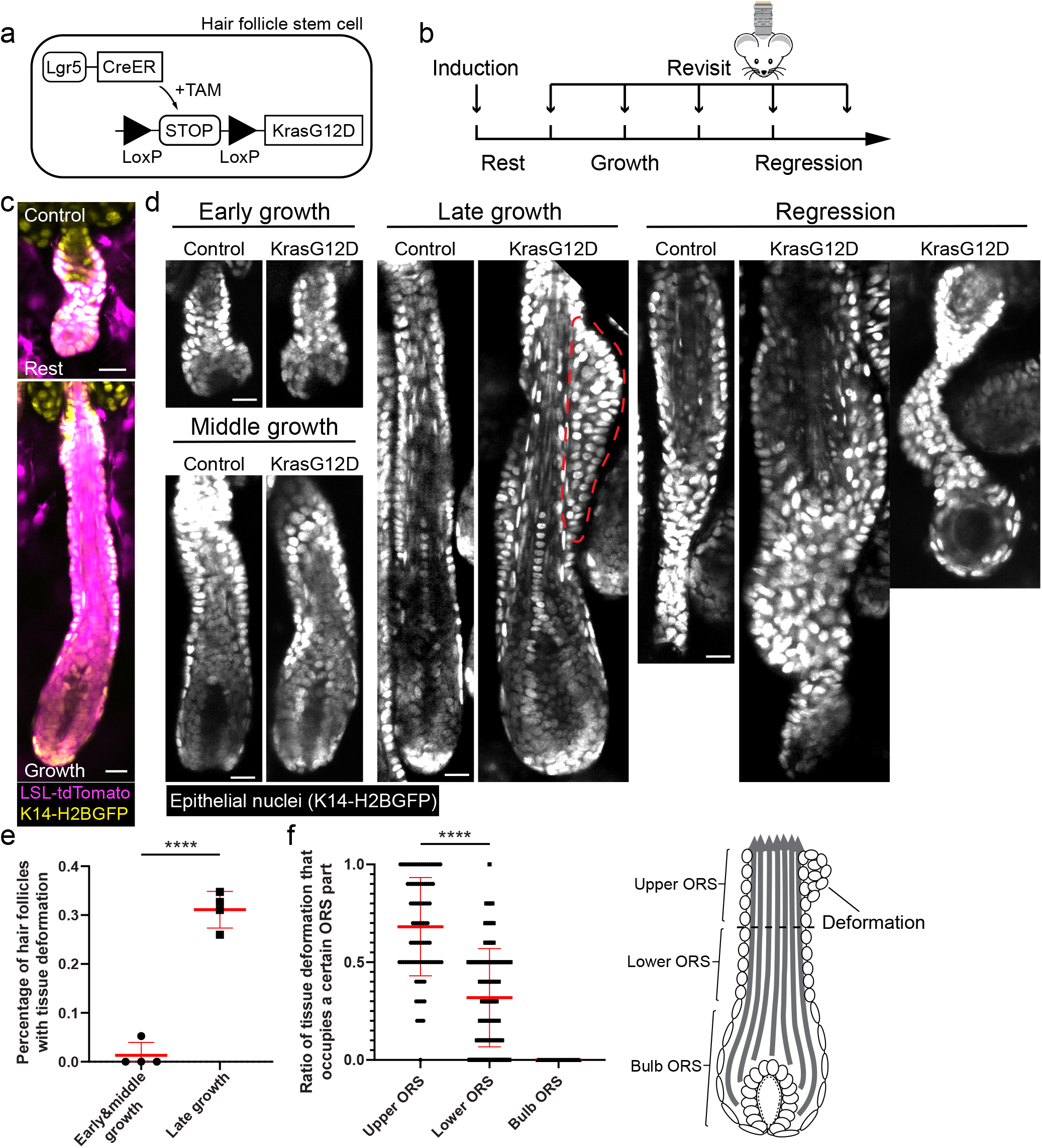
KrasG12D induces spatiotemporally specific tissue deformation in hair follicle regeneration. **a**, Schematic showing the genetic approach to induce KrasG12D in the hair follicle stem cells via tamoxifen (TAM) inducible Cre-LoxP system. **b**, Schematic showing the timing of the KrasG12D induction and repeated imaging relative to the hair cycle stages. **c**, Representative two-photon images of the wild type resting and growing hair follicles carrying Cre-inducible tdTomato (magenta) reporter after induction. **d**, Representative two-photon images of the control and KrasG12D hair follicles at different stages of the hair cycle. Bump-like tissue deformation in the ORS is outlined by a red dashed line. **e**, Percentages of the KrasG12D hair follicles with tissue deformation at different stages of hair follicle growth. n=4 mice. 169 early & middle growth and 311 late growth hair follicles were analyzed. ****, p<0.0001. **f**, Percentages of tissue deformations occupying upper, lower, and bulb ORS for individual KrasG12D hair follicles. 97 KrasG12D hair follicles in panel **e** that developed deformation were analyzed. ****, p<0.0001. No tissue deformation that occupied bulb ORS was detected. Schematic shows different parts of ORS. Two-sided unpaired t-test was used to calculate p values. Epithelial nuclei were labeled by K14-H2BGFP (yellow in **c** and white in **d**). Scale bars, 20 μm.

Through tracking the hair follicles in the same mice, we found that mutant hair follicles did not exhibit obvious architectural defects during the early and middle growth stages (Anagen I-III) (Figure 1d and 1e). However, as hair follicles grew into late stages (started from Anagen IV), bump-like tissue deformations specifically formed in the outermost layer (ORS) of the mutant hair follicles, where the expanded stem cells (ORS cells) are located (Figure 1d and 1e). Consistent with previous studies, the inner layers that consist of hair progenitors and differentiating cells did not show abnormal architecture^5, 8^. When mutant hair follicles entered the regression phase, new architectural abnormalities emerged, including malformed epithelial strand and follicular cyst (Figure 1d, Regression). Importantly, the hair follicles with a delayed entrance to the growth cycle did not show deformation at early growth stages, at the same time as other follicles in the same skin developed the phenotype in the late growth stages. This indicates that the temporal specificity of the Kras-induced tissue abnormality is due to changes between different hair follicle stages, not the duration of exposure to oncogenic Kras (Supplementary figure 2). In addition to the temporal specificity of this tissue aberrancy, we also found that the bump-like tissue deformations preferentially formed at specific positions along the hair follicle length. In the late growth stages, the upper ORS was significantly more likely to form the tissue deformation than the lower ORS, while the bottom ORS at the surface of the hair bulb was never deformed (Figure 1f). Together, these findings show that oncogenic KrasG12D induces tissue deformation in a spatiotemporally specific manner during hair follicle regeneration, particularly a bump-like structure in the upper ORS at the late growth stage.

### KrasG12D causes abnormal cell division and migration during hair follicle growth

The spatiotemporal specificity of the bump-like tissue deformation made us wonder what cellular alterations induce this tissue architecture disruption, and particularly how these alterations vary depending on the different niches within the hair follicle. Previous studies identified high numbers of proliferative cells at relatively late stages of hair follicles with KrasG12D-induced deformation^5, 8^. However, it is still unclear when hyperproliferation or other abnormal cell behaviors initiate relative to tissue deformation, and where these changes occur within the hair follicle. In order to better understand the cellular mechanisms that drive tissue deformation, we performed proliferation analyses in early bump formation stages as well as used time lapses to discover additional abnormal cell behaviors. By using an EdU cell proliferation assay, we found that more proliferative cells were detected in the ORS of the KrasG12D mutant hair follicles (Figure 2a and 2b). Interestingly, in the lower ORS, where the vast majority of proliferation occurs in wild type hair follicles^16^, the number of the EdU+ cells was not different between the control and mutant (Figure 2c). In contrast, when we looked at the relative proportion of proliferation in the upper versus lower ORS, we found it was significantly higher in the Kras mutant hair follicles (Figure 2d). These data suggest that instead of elevating proliferation over the entire ORS, the KrasG12D mutation specifically induces cell division in the upper ORS.

**Figure 2.**
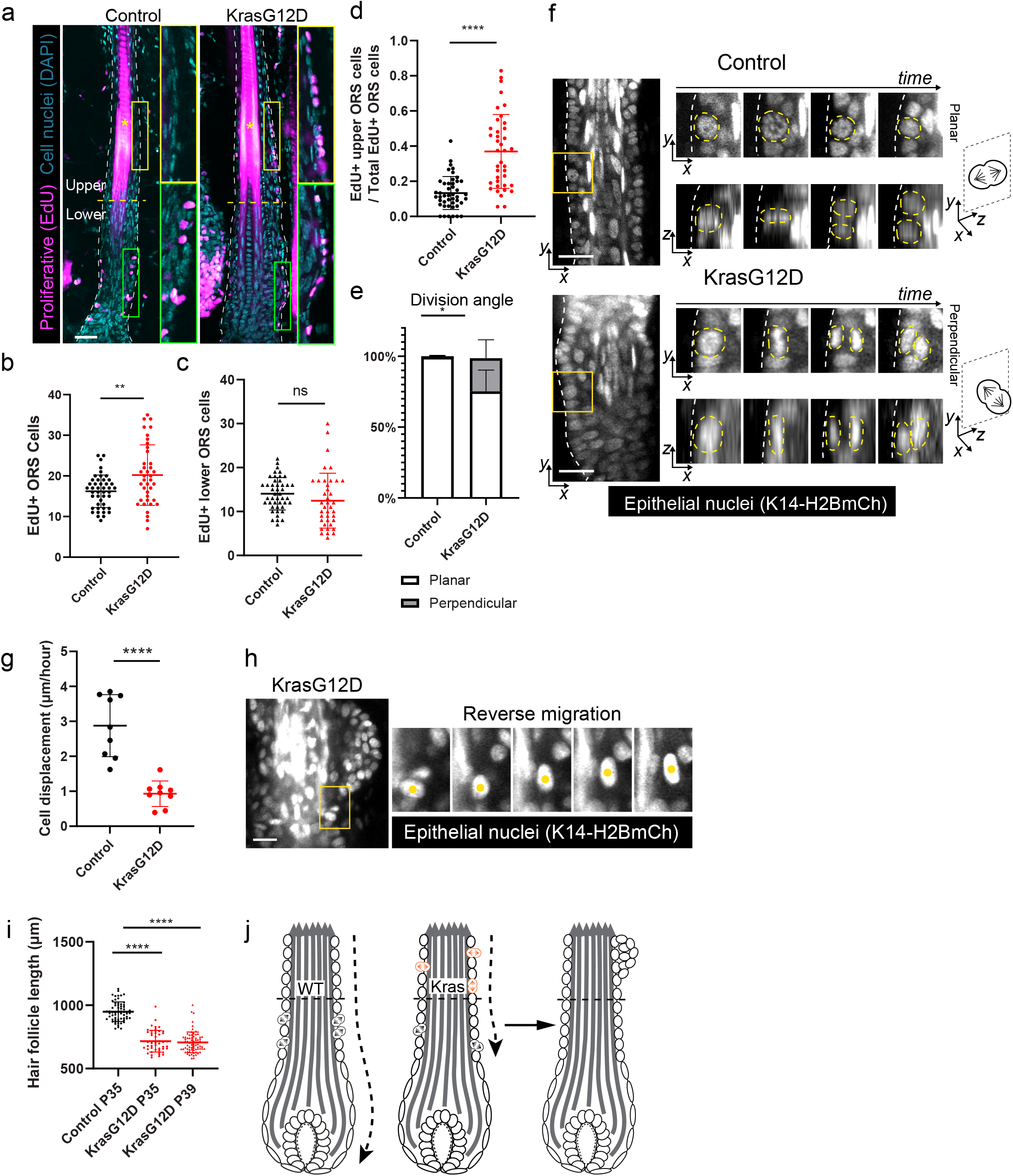
KrasG12D leads to ectopic cell division and migration during hair follicle growth. **a**, Representative images of the control and KrasG12D hair follicles on tissue sections with EdU (magenta) and DAPI (cyan) labeling for analyzing cell proliferation in the ORS. Regions of the ORS analyzed are marked with white dashed lines. Upper and lower ORS are separated by yellow dashed line. Insets show the enlarged regions of the upper and lower ORS. Asterisk, hair shaft. **b**, Total numbers of EdU+ ORS cells in the control and KrasG12D hair follicles. n=45 hair follicles in 4 control mice and 39 hair follicles in 4 KrasG12D mice. **, p=0.0026. **c**, Numbers of EdU+ lower ORS cells in the control and KrasG12D hair follicles. The same hair follicles in **b** were analyzed. ns, not significant, p=0.1550. **d**, Ratio of the EdU+ upper ORS cells to the total EdU+ ORS cells in the control and KrasG12D hair follicles. The same hair follicles in **b** were analyzed. ****, p<0.0001. **e**, Percentages of the planar and perpendicular divisions in the control and KrasG12D hair follicles based on time lapse analysis. n=3 control and 3 KrasG12D mice. 127 divisions in 48 control hair follicles and 147 divisions in 48 KrasG12D hair follicles were analyzed. *, p=0.0378. **f**, Representative two-photon time lapse frames of the control and KrasG12D hair follicles showing division angles of the ORS cells. Basement membranes are marked with white dashed lines. Nuclei of dividing cells are marked with yellow dashed lines. **g**, Average displacement distance per hour of the control and KrasG12D ORS cells. n=9 hair follicles in 3 control mice and 9 hair follicles in 3 KrasG12D mice. ****, p<0.0001. **h**, Two-photon time lapse frames of one representative KrasG12D late growth hair follicle showing one ORS cell migrating in the opposite direction compared to typical downward migration of ORS cells. **i**, Length comparison between the P35 control hair follicles and the P35 and P39 KrasG12D hair follicles. n=61 hair follicles in 3 P35 control mice, 48 hair follicles in 3 P35 KrasG12D mice and 73 hair follicles in 3 P39 KrasG12D mice. ****, p<0.0001. **j**, Schematic showing abnormal division and migration of the KrasG21D ORS cells contribute to tissue deformation. Two-sided unpaired t-test was used to calculate p values. Epithelial nuclei were labeled by K14-H2BGFP (white in **f** and **g**). Scale bars, 20 μm.

Cell proliferation can affect overall tissue architecture in multiple ways, depending on how it is carried out in 3D space. Through time lapse analysis of the 3D hair follicles in live mice, we found that the division angle of the ORS cells was significantly more likely to be perpendicular to the basement membrane in the mutant hair follicles than in the control, where divisions are mostly planar (Figure 2e and 2f). Those perpendicular divisions resulted in one daughter being placed into the suprabasal layer rather than remaining basal, which can contribute to bump formation. This result is reminiscent of the effect of active Kras on footpad epidermis, the skin epithelium without hair follicles^17^.

In addition to the ectopic divisions, we also identified abnormal migrations of the Kras mutant ORS cells via long time lapses (5-8 hours). In wild-type hair follicles at the growth phase, the ORS cells migrate downwards to extend the hair follicle length^13, 15^. However, compared with the control, the Kras mutant ORS cells had significantly shorter displacement distances along the growth axis across multiple hours (Figure 2g). Occasionally, the mutant cells could also completely change their migration direction (Figure 2h). These migration defects together with the abnormal division angles can cause cell accumulation in the upper ORS and ultimately less hair follicle extension. Consistent with that, the Kras mutant hair follicles were significantly shorter that the control across different stages (Figure 2i). Even the mutant hair follicles that were close to the end of the growth phase (P39) remained shorter than the earlier stage (P35) control ones, suggesting the length difference was not due to the delay of hair cycle. Altogether, our findings suggest that the spatially mis-regulated cell division, abnormal division angle and migration caused by KrasG12D mutation may lead to the bump-like tissue deformation (Figure 2j).

### KrasG12D converts pulsatile into sustained ERK activation in the expanded hair follicle stem cells

Ras proteins are core components of multiple signaling pathways. The mis-regulated KrasG12D mutant cell behaviors could be a result of the miscommunication between the cells and their microenvironments. A previous study found ERK activation was upregulated in the deformed Kras mutant hair follicles^8^, raising the question of whether mis-regulated ERK signaling mediates the KrasG12D-induced tissue deformation. Interestingly, recent studies showed that the particular temporal patterns of ERK activation can coordinate collective cell behaviors at the tissue level^18–22^. Thus, in order to assess how KrasG12D changes ERK signal dynamics during hair follicle growth, we used a reporter mouse expressing the ERK-KTR biosensor^23, 24^. This biosensor allows us to capture real time ERK activity measurements at the single cell level in live mice based on its nucleocytoplasmic distribution (Figure 3a, nuclear localization indicates inactive ERK and nuclear exclusion indicates highly active ERK). We focused on analyzing the hair follicle growth stage when ORS deformation starts to emerge (Anagen IV). At this stage, hair follicle epithelial cells form longitudinal layers that are symmetrically organized, including the inner layers of multiple types of differentiated cells and the outermost layer of expanded stem cells (ORS cells). Time lapses of the wild type hair follicles that express the ERK reporter showed that different cell types have distinct ERK activation dynamics (Figure 3b). Specifically, the innermost differentiated cell types show sustained low activation; the middle layers of differentiated cell types show sustained high activation; and the outer layers of cell types including the ORS cells show pulsatile activation. This pattern suggests that different ERK signal dynamics may be associated with cell type-specific behaviors and functions. Notably, the pulsatile ERK activation in the ORS was often manifested as directional wave-like signal spread between cells (Supplementary figure 3a). This resembles the epidermal growth factor receptor (EGFR)-mediated ERK signal propagation between epithelial cells both in culture and in skin epidermis, which was found to direct collective cell migration^19, 21–22^. Given that KrasG12D induces tissue deformation and abnormal cell behaviors including the migration defects specifically in the ORS, we focused on analyzing the ERK signal dynamics in those expanded stem cells.

**Figure 3.**
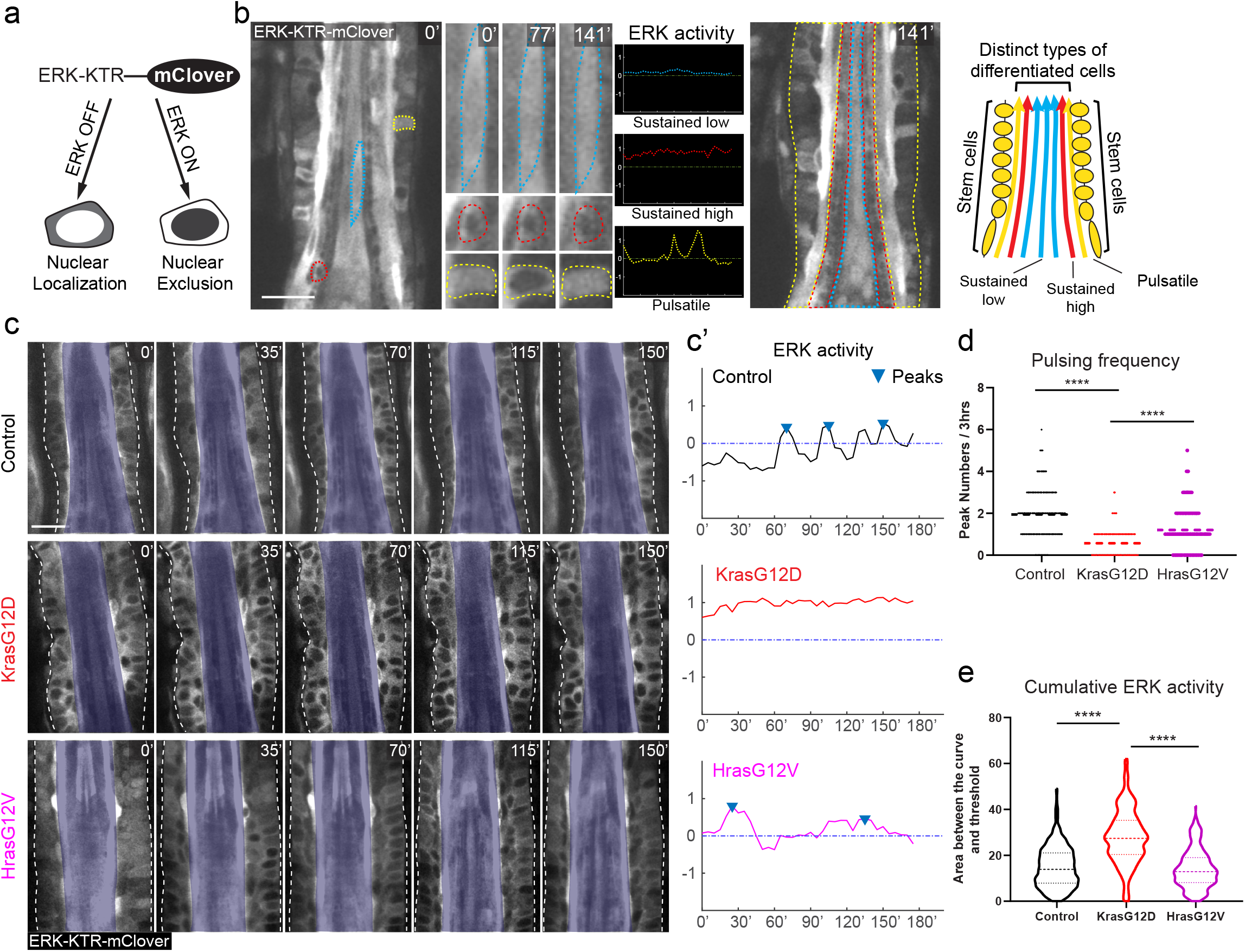
KrasG12D converts pulsatile into sustained ERK activation in the expanded hair follicle stem cells. **a**, Schematic showing that the ERK-KTR-mClover biosensor reports ERK activation status based on its nucleocytoplasmic distribution. **b**, Representative two-photon time lapse frames of the wild type late growth hair follicle expressing the ERK biosensor. Example cells in different layers are outlined by dashed lines (blue, inner differentiated cell; red, middle differentiated cell; yellow, outer stem cell). Their ERK activation dynamics are indicated by the selected time frames and plotted curves. Schematic shows distinct ERK activation dynamics at different layers of the late growth hair follicle. **c**, Representative two-photon time lapse frames of the wild type, KrasG12D, and HrasG12V hair follicles, showing different ERK activation dynamics in the ORS. Hair follicle borders are marked with white dashed lines and inner layers are masked in opaque blue to better show the ORS cells. **c’**, ERK activation curves of one representative ORS cell in each hair follicle in **c**. Peaks within the curves were identified by a threshold of local maxima to reflect the pulsing frequency. **d**, Pulsing frequency of the ERK signal in the wild type, KrasG12D, and HrasG12V ORS cells represented by peak numbers in the 3 hour ERK activation curve. n=448 cells in 32 hair follicles in 3 control mice, 364 cells in 26 hair follicles in 3 KrasG12D mice, and 364 cells in 26 hair follicles in 3 HrasG12V mice. ****, p<0.0001. **e**, Cumulative ERK activity of the wild type, KrasG12D and HrasG12V ORS cells represented by the area between the curve and the threshold. The same cells in **d** were analyzed. ****, p<0.0001. Two-sided unpaired t-test was used to calculate p values. Scale bars, 20 μm.

Time lapses of the Kras mutant hair follicles showed that most ORS cells persistently maintained nuclear exclusion phenotype of the reporter protein, indicating active ERK signaling, in contrast to the frequent switching between nuclear localization and exclusion in the control (Figure 3c, Control and KrasG12D) (Supplementary movie 1 and 2). Further quantitative analysis of the ERK activation in single cells based on the ratio of the cytoplasm-versus nucleus-localized reporter showed that Kras mutant ORS cells maintained sustained ERK activation instead of the pulsatile signals observed in the control (Figure 3c’, Control and KrasG12D, one representative cell for each). By identifying and counting peak numbers in the activation curve, we observed that the KrasG12D ORS cells had significantly lower pulsing frequency when compared with the control (Figure 3d). Furthermore, we found that the Kras mutant ORS cells had much more cumulative ERK signal activation estimated by the area above the activation threshold (Figure 3e). These together suggest that KrasG12D converts pulsatile into sustained ERK activation in the expanded hair follicle stem cells in the late growth stage, which results in high cumulative ERK signal activation.

To further test whether this ERK activation change is associated with the formation of ORS deformation, we analyzed ERK signal dynamics in the hair follicles carrying another oncogenic Ras mutation - HrasG12V, which could potentially alter ERK signal but which we previously showed does not induce bump-like ORS deformation^10^. Using a similar genetic approach as for KrasG12D to induce HrasG12V (Lgr5-CreER; LoxP-Hras-LoxP-HrasG12V) in the ORS cells did not cause sustained ERK activation. Instead, HrasG12V cells had significantly higher pulsing frequency and lower cumulative ERK activity when compared with KrasG12D (Figure 3c-3e, HrasG12V) (Supplementary movie 3). These data suggest that the bump-like tissue deformation in the Kras mutant hair follicles correlates with the sustained ERK activation specifically induced by KrasG12D.

Since the tissue deformation and abnormal cell behaviors in the Kras mutant hair follicles showed spatial specificity, we wondered whether the ERK signal change induced by KrasG12D was also spatially different across regions of the hair follicle. To address this question, we analyzed ERK signal dynamics based on the different positions of the mutant ORS cells and found that upper and lower ORS cells had similar sustained and high ERK signaling levels (Supplementary figure 3b and 3c). These data suggest that the spatial specificity of the tissue abnormality in the Kras mutant hair follicle is caused by spatially distinct factors either downstream of or parallel to the ERK kinase.

Altogether, our data demonstrate that KrasG12D converts pulsatile into sustained ERK activation in the ORS of hair follicles at the late growth stage, which may cause the bump-like tissue deformation.

### Interrupting sustained ERK activation can both prevent and reverse KrasG12D-induced tissue deformation

In order to further understand whether sustained ERK activation induced by KrasG12D was the cause of the tissue deformation in the growing hair follicle, we intradermally injected MEK inhibitor (MEKi) to transiently interrupt the sustained ERK signal in the Kras mutant hair follicles. By using the live ERK reporter, we confirmed that MEKi injection can block the ERK activation in all the skin cells, including the pulsatile ERK activation in the ORS cells of the late growth hair follicles (Figure 4a). However, the effect of the drug can only last for a few hours. The ERK activation reappears 6 hours after MEKi injection (Figure 4b). Therefore, our approach represents a transient blockage of the ERK activity, which we leveraged to interrupt the sustained ERK signal in the Kras mutant cells instead of fully blocking it.

**Figure 4.**
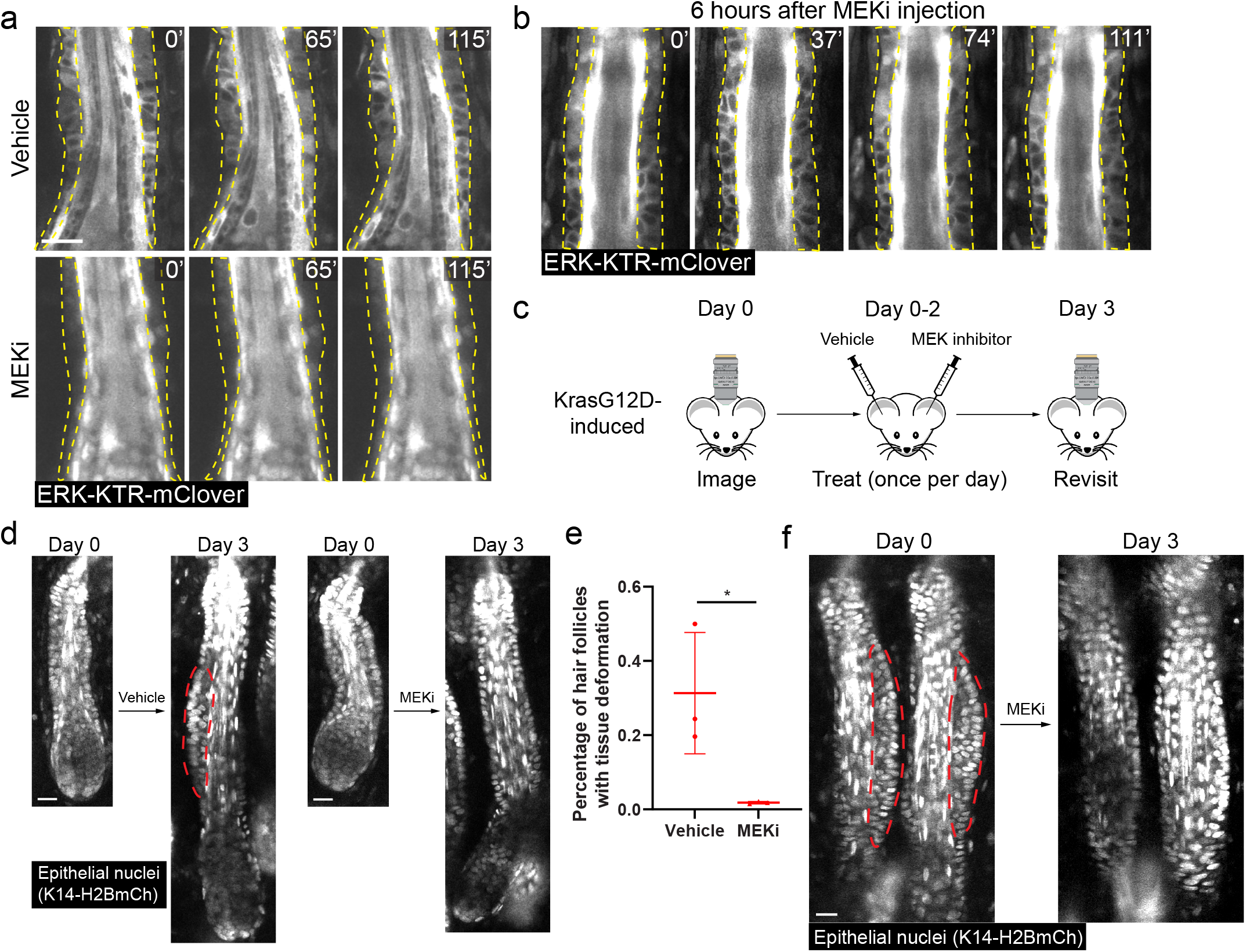
Interrupting sustained ERK activation can both prevent and reverse KrasG12D-induced tissue deformation. **a**, Representative two-photon time lapse frames of the wild type late growth hair follicles expressing the ERK biosensor 2.5 hours after intradermal injection of the MEK inhibitor (MEKi) or vehicle. **b**, Representative two-photon time lapse frames of the wild type late growth hair follicles expressing the ERK biosensor 6 hours after intradermal injection of MEKi. **c**, Schematic showing the schedule of imaging and drug treatment of the same mice for testing the consequence of interrupting sustained ERK activation in the KrasG12D hair follicles. MEKi and vehicle were intradermally injected into the left ear and right ear, respectively, of the same mouse. **d**, Representative two-photon images of the same KrasG12D hair follicles either treated with MEKi or vehicle in the same mouse. **e**, Percentages of the KrasG12D hair follicles having tissue deformation after the treatment of MEKi or vehicle. n=3 mice. 104 vehicle-treated and 170 MEKi-treated hair follicles were analyzed. *, p=0.0352 **f**, Representative two-photon images of the same KrasG12D hair follicles before and after MEKi treatment showing reversal of the tissue deformation. ORS layers are outlined by dashed lines in **a** and **b**. Bump-like tissue deformation in the ORS is outlined by red dashed lines in **d** and **f**. Two-sided unpaired t-test was used to calculate p value. Scale bars, 20 μm.

To assess whether interrupting sustained ERK activation can affect the ORS deformation phenotype in the Kras mutant hair follicles, we combined the live imaging-based longitudinal tracking with the drug injection approach (Figure 4c). We first imaged the hair follicles at the middle and late growth stages on both ears of the Kras mutant mice. Then we intradermally injected MEKi into one ear and vehicle into the other ear as the internal control, once a day for three days. Afterwards, we imaged the same hair follicles again to assess the phenotype. Through this approach, we found that although in the vehicle-injected ear, Kras mutant hair follicles still formed ORS deformation, the vast majority of the hair follicles in the MEKi-injected ear did not show any architecture abnormality (Figure 4a and 4e). More interestingly, for the hair follicles that had already formed tissue deformation prior to the drug treatment, that phenotype was reversed after the three-day MEKi treatment (Figure 4f). Together, these data suggest that the sustained ERK activation induced by KrasG12D is required for both the formation and maintenance of the ORS deformation of the mutant hair follicles.

## Discussion

Cancer formation is a context-dependent process^2^. The same oncogenic driver mutation can cause cancer in one tissue type but produce no phenotype in another. Tumor microenvironment, or ‘niche’, is one of the key factors that cause the tissue-specific differences^1^. Identifying specific niches that support or inhibit oncogenesis and understanding the underlying mechanisms can provide clues that inform therapeutic strategies for cancer prevention and treatment. Here, by using intravital imaging to longitudinally track tissues carrying an oncogenic Kras mutation during hair follicle regeneration, we find that the early events of tissue disruption before cancer formation are also niche-dependent. In particular, spatiotemporally specific conditions in the upper part of the late growth hair follicle synergize with the KrasG12D mutation to induce abnormal cell behaviors and bump-like deformation of the expanded stem cell layer. Further analyses of the activation dynamics of the downstream signaling show that KrasG12D converts pulsatile into sustained ERK activation to induce tissue deformation, and interrupting the sustained ERK signal through temporary MEK inhibition can reverse the abnormal tissue architecture. These together provide new insights into the mechanisms of oncogene-induced tissue disruption, in which oncogenic mutations and their associated signaling dynamics synergize with spatiotemporal cues from the microenvironment to impair orchestrated cell arrangement. Given KrasG12D is a driver mutation for many types of cancers across multiple organs^25^, the mechanism we identified can have broad implications in understanding the fundamental principles of cancer initiation.

ERK signal is a multi-functional pathway that often gets ectopically activated in various cancers. In both normal and pathological conditions, ERK signaling controls many types of cell behaviors such as proliferation, migration, differentiation, and survival. Interestingly, previous studies found that this single signaling pathway can execute those specific function through different kinds of activation dynamics. For example, studies in cell culture and fly embryo showed that the duration of the ERK activation can control proliferation and differentiation decisions^26, 27^. Furthermore, wave-like intercellular ERK signal propagation has been recently discovered in multiple mammalian epithelia, zebrafish scale and fly notum, and was shown to function to coordinate collective cell migration, tissue growth, epithelial sealing, stem cell compartment size control, etc, depending on the specific context^19–22, 28–30^. Here, by using a live ERK signaling reporter, we find that different ERK signaling dynamics are also associated with distinct cell behaviors in the late growth hair follicles. Cells that undergo differentiation exhibit sustained ERK activation, while stem cells that proliferate and collectively migrate downwards show pulsatile activity including wavelike regional signal propagation (Figure 3b and Supplementary figure 3a). We also find the KrasG12D mutant stem cells that undergo ectopic division and migration show abnormal sustained ERK activation. These observations suggest that different ERK activation dynamics may play roles in regulating specific cell behaviors in the hair follicle epithelium. The pulsatile ERK signal in the stem cell layer might be critical for coordinating cell proliferation and migration during hair follicle growth. The sustained ERK activation induced by KrasG12D can disrupt both the intracellular ERK fluctuation and the intercellular signal propagation and ultimately lead to tissue deformation. This disruption depend on long term sustained ERK activity, as even temporary inhibition of the ERK activation (a few hours per day) can reverse the deformation phenotype (Figure 4).

The ERK activation dynamics have been shown *in vitro* to arise from feedback mechanisms at different levels of the signaling pathway^31^. Interestingly, a recent study showed that Dual Specificity Phosphatase 6 (DUSP6), an ERK specific phosphatase, can modulate the level of ERK signal pulses in the skin epidermal cells to initiate their differentiation^32^. Hair follicle stem cells may use the similar feedback network to achieve pulsatile ERK activation inside the cells. KrasG12D might induce sustained ERK signal by bypassing or suppressing those feedback mechanisms via processes that HrasG12V does not affect. This difference between KrasG12D and HrasG12V leads to distinct tissue architecture consequences: both increase cell proliferation in the growing hair follicle (Figure 2b)^10^, but only KrasG12D can induce bump-like tissue deformation. Similar differential effects between Ras oncogenes were observed in other contexts, suggesting Ras isoform-specific mechanisms driving tissue disruption (Li, 2018).

We showed spatially specific abnormal cell proliferation and tissue deformation in the Kras mutant hair follicles at late growth stages (Figure 1 and 2), yet the ERK activation dynamics were changed throughout the hair follicle (Supplementary figure 3b and 3c). These results suggest that additional factors either downstream of or in synergy with ERK account for the spatial specificity of the mutant phenotype. Interestingly, proliferation of the ORS cells in wild type hair follicles is spatially patterned along the axis of growth (Figure 2a and 2d)^16^ and single cell transcriptome analysis recently revealed positionally different transcriptional signatures within the ORS population^12^. More thoroughly characterizing what distinguishes these location-specific cell phenotypes and how they are established in normal hair follicles will be critical for understanding the niche-dependent responses to oncogenic mutations that we have discovered here. Furthermore, as hair follicles undergo dynamic tissue change during regeneration, those spatially patterned compartments need constant readjustment. Since ERK signal propagation has been shown to regulate patterned cell behaviors^22, 29^, the intercellularly coordinated pulsatile ERK activation in the ORS might be necessary to maintain the domains of distinct cell behaviors. Once KrasG12D converts pulsatile into sustained ERK signal, the mutant cells might lose their sense of positions and cause local accumulation in those permissive positions, eventually as manifesting as tissue deformation. Further understanding the functions of the coordinated ERK activation and its molecular mechanisms will shed light on how Ras mutations cause spatially specific tissue abnormalities.

## Methods

### Mice

K14-H2BGFP^33^ mice were obtained from E. Fuchs. Lgr5-IRES-CreER mice were obtained from H. Clevers. K14-H2BmCherry^34^ and ERK-KTR-mClover^23^ mice were generated and described previously. R26-LoxP-STOP-LoxP-tdTomato^35^, LoxP-STOP-LoxP-KrasG12D^36^ and LoxP-Hras-LoxP-HrasG12V^37^ mice were obtained from The Jackson Laboratory. Mice were bred to a mixed albino background. K14-H2BmCherry was always bred together with ERK-KTR-mClover to facilitate nuclei identification.

To induce KrasG12D in the hair follicle stem cells, Lgr5-IRES-CreER; LoxP-STOP-LoxP-KrasG12D (with or without R26-LoxP-STOP-LoxP-tdTomato; K14-H2BGFP or K14-H2BmCherry; ERK-KTR-mClover) mice were given a single dose of tamoxifen (2mg in corn oil) around postnatal day 18 (P18) by intraperitoneal injection. Sibling mice without KrasG12D were used as control. Live imaging for revisiting the same skin was performed every 3-4 days between P25 and P52 in order to capture different stages of hair cycle. Hair follicle stages were determined based on previous literature^38^.

For MEKi treatment, 50 μM MEKi (PD0325901) in PBS with 0.5% DMSO was injected intradermally into one ear of the mice once per day for three days. Vehicle (PBS with 0.5% DMSO) was injected into the other ear of the mice at the same time. Live imaging was performed right before the first injection and one day after the last injection.

Mice from experimental and control groups were randomly selected from either sex for experiments. No blinding was done. All studies and procedures involving animal subjects were approved by the Institutional Animal Care and Use Committee at Yale School of Medicine and conducted in accordance with the approved animal handling protocol.

### Intravital imaging

Imaging procedures were similar to those previously described^39^. Mice were anaesthetized with vaporized isoflurane delivered by a nose cone (1.5% in oxygen and air). Mice were placed on a warming pad during imaging. The ear was mounted on a custom-made stage and a glass coverslip was placed directly against it. A LaVision TriM Scope II (LaVision Biotec) microscope equipped with a Chameleon Vision II (Coherent) two-photon laser (using 940 nm for exciting green fluorophore) and a Chameleon Discovery (Coherent) two-photon laser (using 1120 nm for exciting red fluorophore) was used to acquire z-stacks of 50-200 μm in 3μm steps through either a Nikon 25x/1.10 or a Nikon 40x/1.15 water immersion objective. Optical sections were scanned with a field of view of 0.08 or 0.20 mm^2^. For imaging large areas, multiple tiles (up to 56) of optical fields were captured using a motorized stage. Patterns of hair follicle clusters were used as landmarks for revisiting the same skin area. For capturing real time cell behaviors and ERK signaling dynamics, serial optical stacks were obtained at 8-9 and 5 minutes intervals respectively to generate time lapses.

### EdU cell proliferation assay

Mice were intraperitoneally injected once with 50 μg/g BrdU, 6 hours before tissue collection. Back skins were then dissected and embedded in optimal cutting temperature (OCT; Tissue Tek). Frozen OCT blocks were sectioned at 50 μm to include better 3D structure of the hair follicles. EdU labelling was done by using the Click-iT Alexa Fluor 555 kit (Thermofisher) according to the manufacturer’s instructions. DAPI (Thermofisher) was used for nuclear counterstain afterwards. Large areas of 3D image stacks were then acquired by the LaVision TriM Scope II (LaVision Biotec) microscope.

### Image analysis

Raw image stacks were imported into Fiji (ImageJ, NIH) or Imaris (BitPlane) for analysis. Tiles of optical fields were stitched in Fiji. Selected optical planes or z-projections of sequential optical sections were used to assemble figures.

For quantifying EdU+ cells and hair follicle length, only the hair follicles whose entire length could be captured in the thick sections were selected for analyses. Hair follicle length was measured from the bottom of the bulge to the bottom of the hair follicles to reflect the growth of the cycling portion. Upper and lower ORS were divided by the midline of the hair follicle length above the bulb. Only the EdU+ cells in the middle plane of the hair follicles were quantified in order to normalize the areas of the ORS between hair follicles.

For cell migration analysis, time lapses were imported into Imaris and cell nuclei were identified and tracked by the Spot function. Cell displacement along the hair follicle axis was then calculated based on the distance between nuclei at different time points.

To quantify the ERK signal dynamics, fluorescent intensity of the ERK reporter within the cell nucleus of individual cells at each time point of the time lapse was sample by the Measure function of Fiji. Meanwhile, an average cytoplasmic reporter intensity was sampled similarly for each cell. The log base 2 of the ratio between the background-subtracted cytoplasmic and nuclear intensity was then plotted in Matlab (MathWorks) to generate the activation curve over time. Peak identification within the activation curve was done by the Findpeak function in Matlab with 0.4 Minimum peak prominence. Accumulative ERK activity was calculated by using the Trapz function in Matlab to measure the area between the activation curve and the arbitrary threshold line (−0.3, based on the wild type curves).

### Statistics and reproducibility

Statistical calculations were performed using Prism 9 (GraphPad). An unpaired Student’s t-test was used to determine the significance between two groups. A p value of <0.05 was considered significant; precise p values can be found in the figure legends. No statistical method was used to predetermine sample size. Mouse numbers represent the biological replicates. Sample size and replicates are indicated in the figure legends.

## Data availability

All the data that support the findings of this study are available from the corresponding author upon reasonable request.

## Acknowledgments

We thank the Greco lab members for helpful discussion. We thank Hans Clevers for the Lgr5-IRES-CreER mice and Elaine Fuchs for the K14-H2BGFP mice. This work is supported by the HHMI Scholar award, NIH grants number 1R01AR063663, 5R01AR067755 and 1DP1AG066590, and Dermatology Foundation Research Grant. V.G. was a New York Stem Cell Foundation Robertson Investigator. T.X. was supported by the James Hudson Brown - Alexander Brown Coxe Postdoctoral Fellowship and the New York Stem Cell Foundation Druckenmiller Fellowship.

## Author Contributions

T.X., S.R. and V.G. designed the experiments. T.X. performed the experiments and analyzed the data. S.G. characterized and maintained the KrasG12D and HrasG12V mice. D.G. assisted with the two-photon imaging and data analysis. T.X, L.E.G., S.R. and V.G. wrote the manuscript with input from all the authors.

## Supplementary materials

**Supplementary figure 1.**
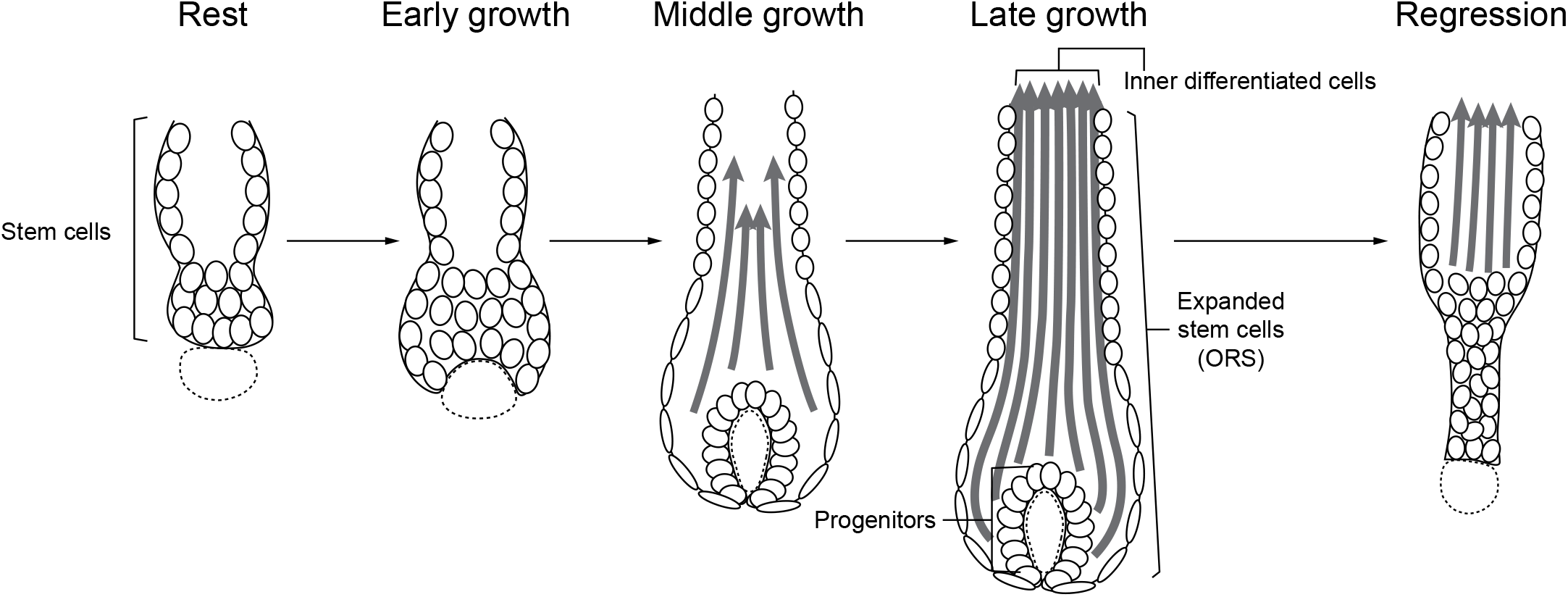
Schematic showing different stages of the hair follicle regeneration cycle.

**Supplementary figure 2.**
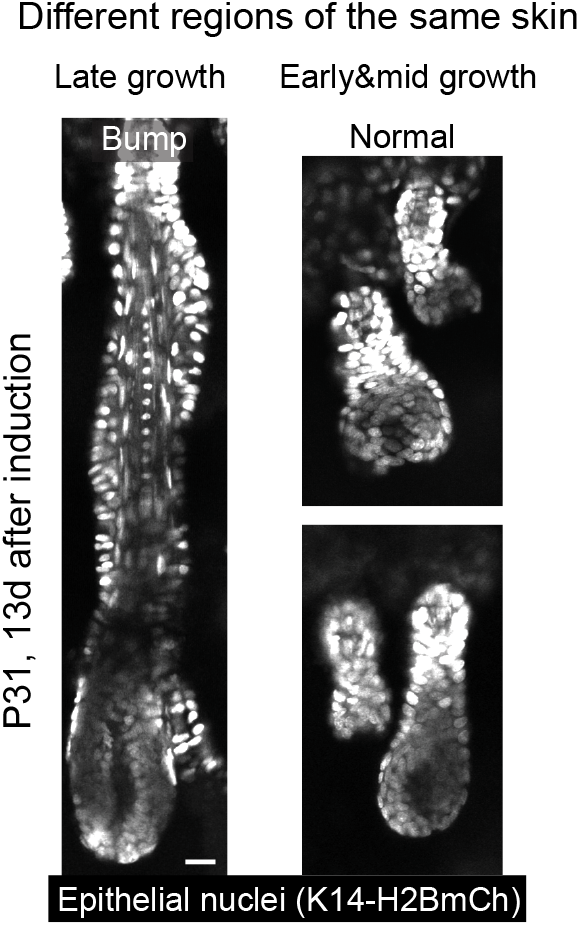
Representative two-photon images of the KrasG12D hair follicles in the same mouse 13 days after induction. In the skin region that hair follicles entered late growth stages, bump-like deformation emerged in the ORS, while in the area of early and middle growth stages, hair follicles were normal.

**Supplementary figure 3.**
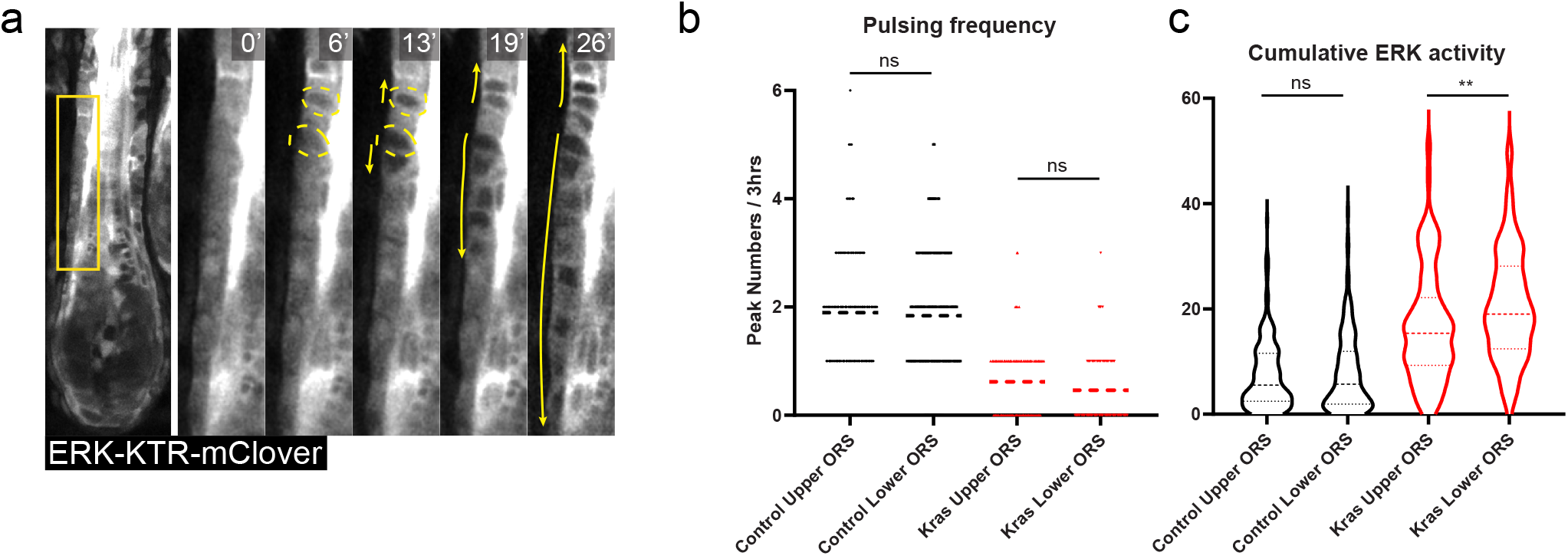
**a**, Representative two-photon time lapse frames of the wild type late growth hair follicle expressing the ERK biosensor showing wave-like ERK signal propagation in the ORS. **b**, Pulsing frequency of the ERK signal in the upper and lower ORS cells of the control and KrasG12D hair follicles. n=236 upper and 162 lower ORS cells in 3 wild type mice, 1666 upper and 91 lower ORS cells in KrasG12D mice. ns, not significant, p=0.6385 and 0.0571. **c**, Accumulative ERK activity of the upper and lower ORS cells of the control and KrasG12D hair follicles. The same cells in **b** were analyzed. ns, not significant, p=0.4481, **, p=0.0079.

**Supplementary movie 1.**

Two-photon time lapse of a representative control late growth hair follicle.

**Supplementary movie 2.**

Two-photon time lapse of a representative KrasG12D late growth hair follicle.

**Supplementary movie 3.**

Two-photon time lapse of a representative HrasG12V late growth hair follicle.

## References

1. Bissell, M.J. & Hines, W.C. Why don’t we get more cancer? A proposed role of the microenvironment in restraining cancer progression. Nat Med 17, 320–329 (2011).

2. Schneider, G.,Schmidt-Supprian, M., Rad, R. & Saur, D. Tissue-specific tumorigenesis: context matters. Nat Rev Cancer 17, 239–253 (2017).

3. Kakiuchi, N. & Ogawa, S. Clonal expansion in non-cancer tissues. Nat Rev Cancer 21, 239–256 (2021).

4. Amberg, N. et al. Mouse models of nonmelanoma skin cancer. Methods Mol Biol 1267, 217–250 (2015).

5. Lapouge, G. et al. Identifying the cellular origin of squamous skin tumors. Proc Natl Acad Sci U S A 108, 7431–7436 (2011).

6. Latil, M. et al. Cell-Type-Specific Chromatin States Differentially Prime Squamous Cell Carcinoma Tumor-Initiating Cells for Epithelial to Mesenchymal Transition. Cell Stem Cell 20, 191–204 e195 (2017).

7. White, A.C. et al. Stem cell quiescence acts as a tumour suppressor in squamous tumours. Nat Cell Biol 16, 99–107 (2014).

8. White, A.C. et al. Defining the origins of Ras/p53-mediated squamous cell carcinoma. Proc Natl Acad Sci U S A 108, 7425–7430 (2011).

9. Brown, S. et al. Correction of aberrant growth preserves tissue homeostasis. Nature 548, 334–337 (2017).

10. Pineda, C.M. et al. Hair follicle regeneration suppresses Ras-driven oncogenic growth. J Cell Biol 218, 3212–3222 (2019).

11. Lee, J. & Tumbar, T. Hairy tale of signaling in hair follicle development and cycling. Semin Cell Dev Biol 23, 906–916 (2012).

12. Joost, S. et al. The Molecular Anatomy of Mouse Skin during Hair Growth and Rest. Cell Stem Cell 26, 441–457 e447 (2020).

13. Rompolas, P. et al. Live imaging of stem cell and progeny behaviour in physiological hair-follicle regeneration. Nature 487, 496–499 (2012).

14. Rompolas, P., Mesa, K.R. & Greco, V. Spatial organization within a niche as a determinant of stem-cell fate. Nature 502, 513–518 (2013).

15. Xin, T., Gonzalez, D., Rompolas, P. & Greco, V. Flexible fate determination ensures robust differentiation in the hair follicle. Nat Cell Biol 20, 1361–1369 (2018).

16. Sequeira, I. & Nicolas, J.F. Redefining the structure of the hair follicle by 3D clonal analysis. Development 139, 3741–3751 (2012).

17. Morrow, A., Underwood, J., Seldin, L., Hinnant, T. & Lechler, T. Regulated spindle orientation buffers tissue growth in the epidermis. Elife 8(2019).

18. Aikin, T.J., Peterson, A.F., Pokrass, M.J., Clark, H.R. & Regot, S. MAPK activity dynamics regulate non-cell autonomous effects of oncogene expression. Elife 9(2020).

19. Aoki, K. et al. Propagating Wave of ERK Activation Orients Collective Cell Migration. Dev Cell 43, 305–317 e305 (2017).

20. De Simone, A. et al. Control of osteoblast regeneration by a train of Erk activity waves. Nature 590, 129–133 (2021).

21. Hino, N. et al. ERK-Mediated Mechanochemical Waves Direct Collective Cell Polarization. Dev Cell 53, 646–660 e648 (2020).

22. Hiratsuka, T. et al. Intercellular propagation of extracellular signal-regulated kinase activation revealed by in vivo imaging of mouse skin. Elife 4, e05178 (2015).

23. Pokrass, M.J. et al. Cell-Cycle-Dependent ERK Signaling Dynamics Direct Fate Specification in the Mammalian Preimplantation Embryo. Dev Cell 55, 328–340 e325 (2020).

24. Regot, S., Hughey, J.J., Bajar, B.T., Carrasco, S. & Covert, M.W. High-sensitivity measurements of multiple kinase activities in live single cells. Cell 157, 1724–1734 (2014).

25. Li, S., Balmain, A. & Counter, C.M. A model for RAS mutation patterns in cancers: finding the sweet spot. Nat Rev Cancer 18, 767–777 (2018).

26. Johnson, H.E. & Toettcher, J.E. Signaling Dynamics Control Cell Fate in the Early Drosophila Embryo. Dev Cell 48, 361–370 e363 (2019).

27. Marshall, C.J. Specificity of receptor tyrosine kinase signaling: transient versus sustained extracellular signal-regulated kinase activation. Cell 80, 179–185 (1995).

28. Gagliardi, P.A. et al. Collective ERK/Akt activity waves orchestrate epithelial homeostasis by driving apoptosis-induced survival. Dev Cell 56, 1712–1726 e1716 (2021).

29. Pond, K.W. et al. Live-cell imaging in human colonic monolayers reveals ERK waves limit the stem cell compartment to maintain epithelial homeostasis. Elife 11, e78837 (2022).

30. Valon, L. et al. Robustness of epithelial sealing is an emerging property of local ERK feedback driven by cell elimination. Dev Cell 56, 1700–1711 e1708 (2021).

31. Dessauges, C. et al. Optogenetic actuator - ERK biosensor circuits identify MAPK network nodes that shape ERK dynamics. Mol Syst Biol 18, e10670 (2022).

32. Hiratsuka, T., Bordeu, I., Pruessner, G. & Watt, F.M. Regulation of ERK basal and pulsatile activity control proliferation and exit from the stem cell compartment in mammalian epidermis. Proc Natl Acad Sci U S A 117, 17796–17807 (2020).

33. Tumbar, T. et al. Defining the epithelial stem cell niche in skin. Science 303, 359–363 (2004).

34. Mesa, K.R. et al. Niche-induced cell death and epithelial phagocytosis regulate hair follicle stem cell pool. Nature 522, 94–97 (2015).

35. Madisen, L. et al. A robust and high-throughput Cre reporting and characterization system for the whole mouse brain. Nat Neurosci 13, 133–140 (2010).

36. Jackson, E.L. et al. Analysis of lung tumor initiation and progression using conditional expression of oncogenic K-ras. Genes Dev 15, 3243–3248 (2001).

37. Chen, X. et al. Endogenous expression of Hras(G12V) induces developmental defects and neoplasms with copy number imbalances of the oncogene. Proc Natl Acad Sci U S A 106, 7979–7984 (2009).

38. Muller-Rover, S. et al. A comprehensive guide for the accurate classification of murine hair follicles in distinct hair cycle stages. J Invest Dermatol 117, 3–15 (2001).

39. Pineda, C.M. et al. Intravital imaging of hair follicle regeneration in the mouse. Nat Protoc 10, 1116–1130 (2015).

